# Linoleic acid metabolite, 13-*S*-hydroxyoctadecadienoic acid, suppresses cancer cell growth by inhibiting mTOR

**DOI:** 10.1101/2025.04.02.646927

**Authors:** Seung Ju Park, Hongmok Kwon, A Yeong Yang, Boah Lee, Haein Lee, Seung Eun Park, Seulgi Lee, Bernie Byunghoon Park, YunHye Kim, Jinwook Lee, Byung-Chul Oh, Daniel Deredge, Patrick L. Wintrode, Mi-Young Kim, Youngjoo Byun, Seyun Kim

**Author notes:** **Correspondence:** (Y.B.), (S.K.). ERSTEQ co., Ltd, Daejeon 34013, Republic of Korea (Seung Ju Park).

## Abstract

Mechanistic target of rapamycin (mTOR) is a key protein kinase that integrates various internal and external signals to control biological events including cell growth. Whereas substantial efforts were made to elucidate protein subunits interacting with mTOR, endogenous metabolite-mTOR interactions remain largely unknown. Using affinity protein purification and mass spectrometry, we identified direct binding of mTOR to 13-*S*-hydroxyoctadecadienoic acid (13-*S*-HODE) which is an oxygenated metabolite of linoleic acid, a polyunsaturated essential fatty acid. Interaction of 13-*S*-HODE with the catalytic ATP-binding domain of mTOR prevented its kinase activity in an ATP-competitive manner. Furthermore, either 13-*S*-HODE treatment or expression of arachidonate 15-lipoxygenase (ALOX15), an enzyme responsible for 13-*S*-HODE production, reduced mTOR signaling, thereby suppressing the growth of cancer cells as well as tumor xenografts. Our results highlight the importance of 13-*S*-HODE serving as a tumor suppressive, mTOR-inhibiting metabolite that links polyunsaturated fatty acid metabolism and the mTOR signaling in controlling cancer cell growth.

## Introduction

Multiple mechanisms have evolved to sense fluctuations in a wide range of cellular small molecules, including nutrients and their modified metabolites, to coordinate signaling networks (Efeyan et al., 2015; Milanesi et al., 2020; Wang and Lei, 2018). Elucidating the molecular interactions among macromolecules and small molecules is essential for fully uncovering signaling networks (Li et al., 2010; Li and Snyder, 2011; Schreiber, 2005). Recently, although small metabolite regulators of proteins have become the significant focus of attention, a limited number of metabolite-protein interactions have been identified relative to other extensive protein-protein and protein-nucleic acid interaction networks (Li et al., 2010; McFedries et al., 2013; Tagore et al., 2008).

Mechanistic target of rapamycin (mTOR) is a key signaling node integrating various environmental signals from growth factors, stress, and nutrients to control major biological events such as growth and metabolism (Ma and Blenis, 2009; Saxton and Sabatini, 2017; Shimobayashi and Hall, 2014). Major efforts to understand mTOR-binding proteins has allowed us to define two multiprotein complexes, mTOR complex 1 (mTORC1) and 2 (mTORC2): mTORC1 controls cellular growth by phosphorylating S6 kinase 1 (S6K1), whereas mTORC2 regulates cell survival by phosphorylating Akt (also known as protein kinase B) and serum/glucocorticoid regulated kinase 1 (SGK1) (Kim et al., 2002; Sarbassov et al., 2004; Sarbassov et al., 2005; Saxton and Sabatini, 2017). It is clear that the dysregulation of the mTOR signaling pathway occurs in many human diseases, including cancer (Guertin and Sabatini, 2007; Mossmann et al., 2018). Importantly, it has been found that majority of human cancers are associated with aberrant activation of mTOR (Cornu et al., 2013; Menon and Manning, 2008; Zhou and Huang, 2011). Thus, dissecting the mTOR-controlling molecular pathways could offer new insights into a better understanding of the mTOR signaling networks and the development of mTOR-targeting therapeutics. Numerous protein factors that physically interact with mTOR and regulate its activity have been identified (e.g., raptor (regulatory-associated protein of mTOR), rictor (rapamycin-insensitive companion of mTOR), DEPTOR (DEP domain-containing mTOR-interacting protein), PRAS40 (proline-rich Akt substrate of 40 kDa) (Kim et al., 2002; Sarbassov et al., 2004; Hara et al., 2002; Peterson et al., 2009; Sancak et al., 2007), whereas phosphatidic acid is the only endogenous metabolite proposed to bind and stimulate mTORC1 (Fang et al., 2001). However, identifying other cellular metabolites that may interact with mTOR and its associated pathways remains to be established.

Herein, we identify 13-*S*-hydroxyoctadecadienoic acid (13-*S*-HODE), an oxidized linoleic acid metabolite, as an mTOR-associated small molecule. Mechanistically, 13-*S*-HODE directly binds to the catalytic domain of mTOR, thereby acting to inhibit mTOR kinase in an ATP-competitive manner. Consequently, 13-*S*-HODE treatment or overexpression of ALOX15 led to the suppressed mTOR signaling, decreased proliferation, and reduced the growth of cultured cancer cells and tumor xenografts. Collectively, our findings reveal that 13-*S*-HODE as a tumor-suppressive polyunsaturated fatty acid (PUFA) metabolite, plays a key role in inhibiting mTOR, thereby suppressing cancer cell growth.

## Results

### Affinity protein purification and mass spectrometry reveals 13-*S*-HODE as an mTOR-associated metabolite

To identify mTOR-binding small metabolites, we performed affinity protein purification followed by liquid chromatography–mass spectrometry (LC-MS) (Figure 1A). FLAG-tagged mTOR was overexpressed in human embryonic kidney 293T (HEK293T) cells and immunopurified. Metabolites extracted from FLAG-mTOR immunoprecipitates were analyzed via LC-MS metabolite profiling based on accurate molecular mass and retention time. A peak ([M-H]^-^: 295.2264 amu) at 6.51 min in the negative ion mode (Figure 1B) was identified as 13-hydroxyoctadecadienoic acid (13-HODE), an oxygenated product of linoleic acid (Spiteller, 1998), the essential polyunsaturated fatty acid (PUFA), on the basis of the same retention time and molecular mass of pure 13-*S*-HODE (Figures 1B and 1C). To verify these findings, HEK293 cell lysates were incubated with either biotin or biotin-labeled 13-*S*-HODE. Notably, biotin-labeled 13-*S*- HODE bound to mTOR and its complex subunit, raptor, but was competed off in the presence of unlabeled 13-*S*-HODE (Figure 1D), clearly supporting associations between 13-*S*-HODE and mTOR in cells.

**Figure 1.**
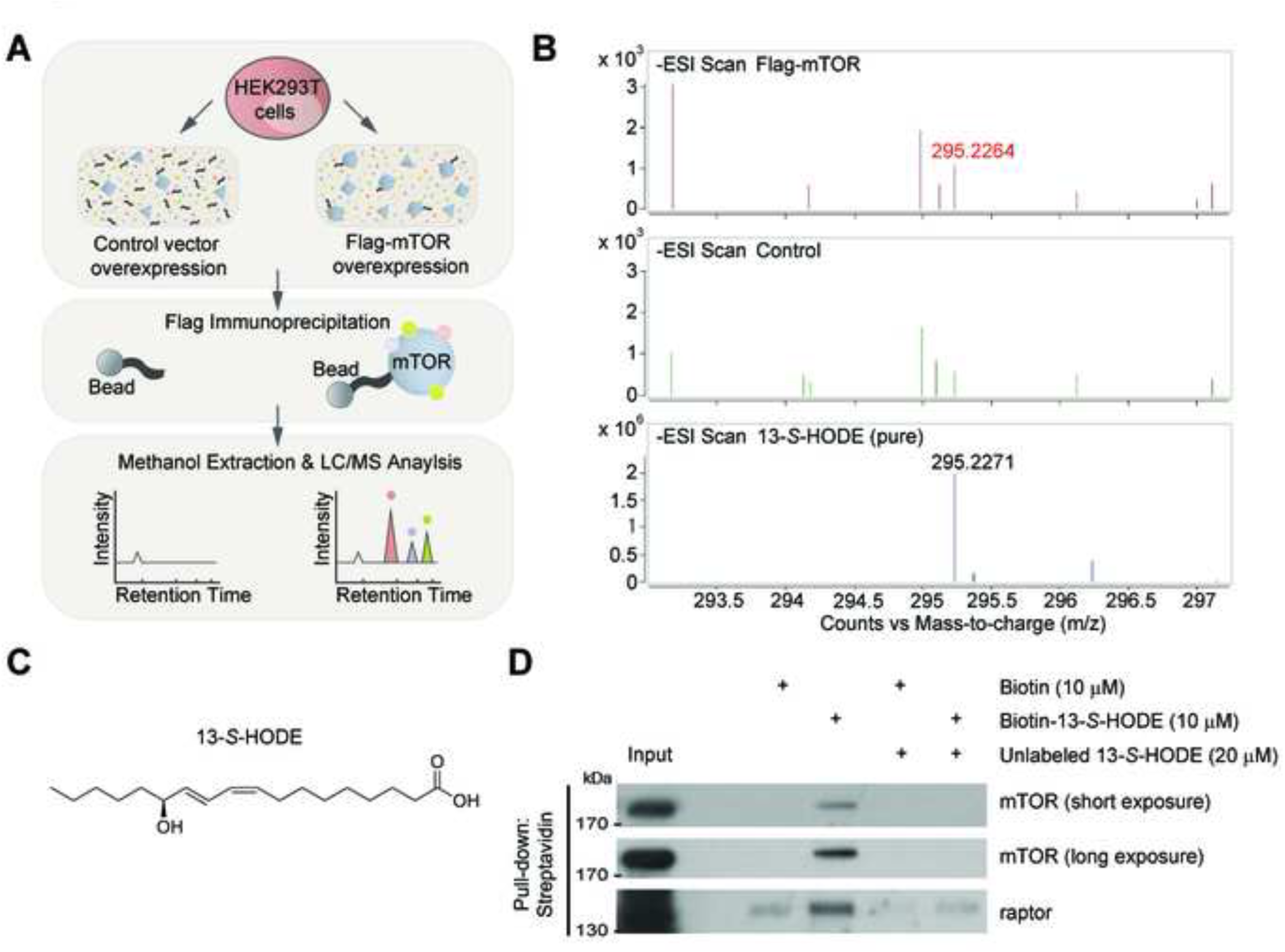
Identification of 13-*S*-HODE as an mTOR-associated metabolite. (A) Experimental schematics for screening endogenous small molecules bound to mTOR. Metabolites specifically bound Flag-mTOR immunoprecipitates relative to control were identified using LC-MS. (B) Extracted ion chromatographs of FLAG-mTOR (top panel) and negative control (middle panel) at 6.51 min in the negative mode. The abundant metabolite ([M-H]^-^: 295.2264) in FLAG-mTOR was confirmed as 13-*S*-HODE by comparison with accurate determinations of the molecular mass and retention times of pure 13-*S*-HODE ([M-H]^-^: 295.2271, lower panel) under the same LC-MS conditions. (C) Chemical structure of 13-*S*-HODE. (D) Binding of biotin-labeled 13-*S*-HODE to endogenous mTOR or raptor in HEK293T cell lysates assessed via streptavidin bead pull-down and immunoblotting. Blots are representative of three independent experiments.

### 13-*S*-HODE directly inhibits mTOR kinase activity in vitro

We examined whether 13-*S*-HODE directly influences mTOR kinase activity to establish the physiological significance of 13-*S*-HODE−mTOR interactions. To address this issue, in vitro kinase assays were performed using either raptor or rictor immunoprecipitates prepared from HEK293T cells. 13-*S*-HODE inhibited phosphorylation at T389 of the mTORC1 substrate, S6K1 in a dose-dependent manner (Figure 2A). Incubation with 13-*S*-HODE additionally blocked phosphorylation of the mTORC2 substrate, Akt, at S473 (Figure 2B), indicating inhibitory activity of the metabolite against both mTORC1 and mTORC2. Relative to the concentration of 13-*S*- HODE required for mTORC1 inhibition, a higher amount was needed to achieve the same level of suppression for mTORC2 (Figures 2A and 2B), suggesting selective activity toward mTORC1.

**Figure 2.**
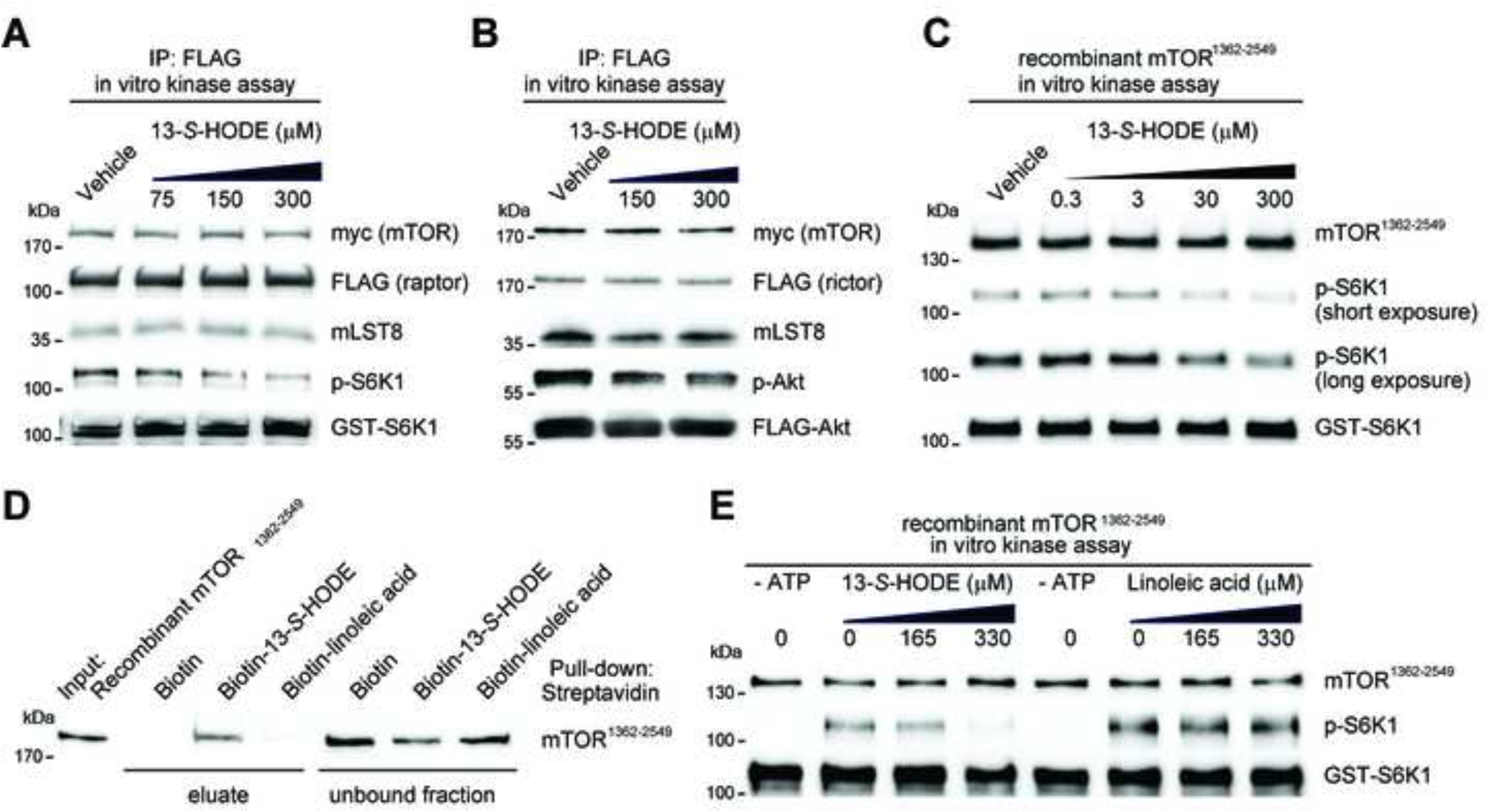
13-*S*-HODE directly inhibits mTOR kinase activity. (A and B) In vitro mTOR kinase assays were performed in the absence or presence of 13-*S*-HODE. FLAG-raptor or FLAG-rictor was cotransfected in HEK293T cells with myc- tagged mTOR. mTORC1 kinase activity was assessed by using FLAG-raptor immunoprecipitates via immunoblotting of T389 S6K1 phosphorylation (A). mTORC2 kinase activity was analyzed by using FLAG-rictor immunoprecipitates, based on S473 Akt phosphorylation (B). (C) In vitro kinase assays were performed using the recombinant human mTOR C- terminal region (mTOR^1362-2549^). mTOR kinase activity was evaluated based on the T389 S6K1 phosphorylation level. (D) Binding of biotin-labeled 13-*S*-HODE to recombinant mTOR^1362-2549^ was assessed via streptavidin bead pull-down and immunoblotting. (E) In vitro kinase assays were performed using the recombinant human mTOR C- terminal region (mTOR^1362-2549^) in the absence or presence of either linoleic acid or 13-*S*-HODE. All panels show representative experiments that were repeated independently three times.

For further confirmation of whether 13-*S*-HODE directly interacts the mTOR kinase domain, we conducted an in vitro kinase assay using the recombinant mTOR C-terminal fragment encompassing amino acid residues 1362 to 2549 that form the intact kinase domain. Notably, 13-*S*-HODE, but not vehicle, inhibited S6K1 phosphorylation (Figure 2C) while failing to exert inhibitory effects on other kinases, including phosphatidylinositol 3-kinase-related kinases (e.g., phosphatidylinositol 3-kinase C3, DNA-dependent protein kinase and extracellular signal–regulated kinase) (Figure S1), highlighting the selective inhibitory activity of 13-*S*-HODE against mTOR. In pulldown experiments, we further detected interactions of biotin-labeled 13-*S*-HODE with the recombinant mTOR C-terminal region (Figure 2D). This binding was specific, since biotin-labeled linoleic acid used as a control did not interact with the mTOR C-terminal fragment nor inhibit its kinase activity (Figures 2D and 2E). Our findings collectively suggest that 13-*S*-HODE inhibits mTOR activity through direct interactions with the kinase domain.

### 13-*S*-HODE binds to the kinase domain of mTOR and acts as an ATP-competitive inhibitor

To gain insights into the mechanism of action of 13-*S*-HODE and establish the potential binding site, we focused on the structural and conformational consequences of 13-*S*-HODE binding to the mTOR C-terminus with the aid of hydrogen/deuterium exchange mass spectrometry (HDX-MS). In the presence of 13-*S*-HODE, several regions exhibited altered deuterium uptake kinetics, which predominantly localized to the kinase domain (Figure S2). A more detailed examination revealed that peptides including residues 1943–1956 at the linker region between the FAT (FRAP, ATM, TRRAP) and FKBP12-rapamycin binding (FRB) domains, 2186–2196, 2270–2280, 2303–2314 or 2421–2431 of the kinase domain of mTOR displayed a significant decrease in exchange at the earliest experimental time-point of D_2_O incubation (10 sec) (Figures S2C and S2D), suggestive of potential ligand-binding sites. Mapping of protected regions on the crystal structure of mTOR (Yang et al., 2013) revealed that the potential 13-*S*-HODE binding site spans across the catalytic cleft of the kinase protein (Figure 3A).

**Figure 3.**
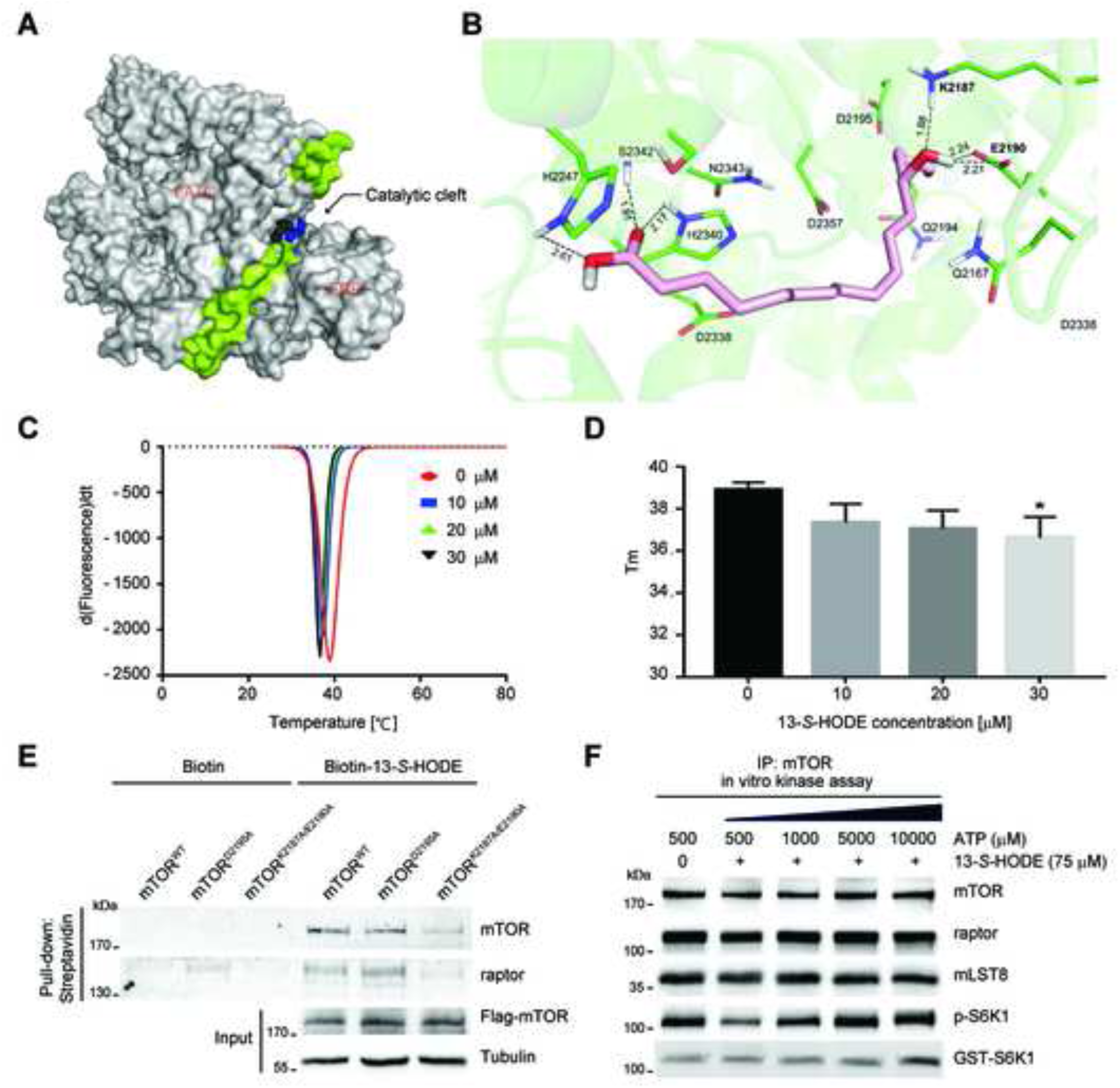
13-*S*-HODE acts as an ATP-competitive inhibitor. (A) Peptides showing significant deviation were mapped onto the structure of mTOR (PDB: 4JSV) on a surface representation. Regions with a significant decrease in deuterium uptake in the presence of 13-*S*-HODE are colored green. Magnesium ions are presented in a black sphere and ADP is labeled in blue to highlight the catalytic site. (B) Optimal docking pose of 13-*S*-HODE in the catalytic site of mTOR kinase. Hydrogen bond interactions between 13-*S*-HODE and mTOR were observed (OH group of 13-HODE with Lys2197 and Glu2190, COOH group with His2340, Ser2342, and His2247). The numbers indicate intermolecular hydrogen bond distance in Å. (C and D) The thermal denaturation of recombinant mTOR kinase domain was measured in the absence and presence of varying lomitapide concentrations, as indicated. Representative derivative (dF/dT) curves are shown for untreated and treated with varying ligand concentrations (C). Midpoint temperatures of the protein-unfolding transition (Tm) are presented as bars (D). Data are means ± SD of at least three independent measurements (\**P* < 0.05). (E) Binding of biotin-labeled 13-*S*-HODE to mTOR was assessed via streptavidin bead pull-down using lysates prepared from HEK293T cells overexpressing FLAG-mTOR^WT^, mTOR^D2195^ or mTOR^K2187A/E2190A^. (F) In vitro mTORC1 activity of raptor immunoprecipitates prepared from HEK293 cells assayed in the presence of 75 μM 13-*S*-HODE and increasing concentrations of ATP. T389 phosphorylation of S6K1 was measured via immunoblotting. Representative experiments are shown, which were repeated independently three times (E and F), all with similar results.

Based on these results, we further performed docking simulations of 13-*S*-HODE at the catalytic site of mTOR kinase to elucidate its potential binding mode. The best docked pose of 13-*S*-HODE was located close to the ATP-binding site of the catalytic cleft of mTOR and formed hydrogen bonds with several residues including Lys2187, Glu2190, His2247, His2340, and Ser2342 (Figure 3B). Two of the residues, Lys2187 and Glu2190, were located in areas potentially protected by 13-*S*-HODE according to HDX-MS data (Figures 3A, S2C, and S2D). These amino acids have also been documented as the key interacting residues with the ADP-MgF_3_-Mg_2_ complex in the mTOR crystal structure (Yang et al., 2013), leading to the proposal that 13-*S*-HODE acts as an ATP-competitive inhibitor.

We next tested direct interaction between 13-*S*-HODE and purified mTOR kinase domain. The thermal stability of recombinant mTOR kinase domain was evaluated by varying concentrations of 13-*S*-HODE. We found that 13-*S*-HODE reduced the thermal stability of recombinant mTOR kinase domain in a concentration-dependent manner, i.e. 30 µM of 13-*S*-HODE decreased the Tm by approximately 2.29 °C (Figures 3C and 3D), suggesting that 13-*S*-HODE directly binds to the kinase domain of mTOR. To clarify whether 13-*S*-HODE–mTOR interactions represent a specific event occurring within the ATP-binding pocket of mTOR, we focused on the residues of mTOR required for interactions with 13-*S*-HODE. Based on docking simulation data, mTOR^D2195A^ and mTOR^K2187A/E2190A^ mutants were generated. Using lysates prepared from HEK293T cells overexpressing wild-type mTOR (mTOR^WT^), mTOR^D2195A^ or mTOR^K2187A/E2190A^ mutants, we performed pull-down assays with biotin-labeled 13-*S*-HODE. Interactions of 13-*S*- HODE with mTOR^K2187A/E2190A^ but not mTOR^D2195A^ were markedly reduced relative to mTOR^WT^ (Figure 3E), demonstrating the specific binding between 13-*S*-HODE and ATP-binding catalytic core of mTOR. The potential ATP-competitive mechanism of 13- *S*-HODE was further investigated by performing an in vitro mTOR kinase assay in the presence of 13-*S*-HODE along with various concentrations of ATP. Notably, the inhibitory effect of 13-*S*-HODE on mTOR was abolished with increasing ATP concentrations (Figure 3F), validating 13-*S*-HODE action through an ATP-competitive mechanism.

### 13-*S*-HODE inhibits mTOR signaling and cancer cell growth

In multiple human cancer types, nearly undetectable levels of 13-*S*-HODE and arachidonate 15-lipoxygenase (*ALOX15*), a rate-limiting enzyme that synthesizes 13-*S*- HODE from linoleic acid have been reported (Lee et al., 2011; Shureiqi et al., 2003; Shureiqi et al., 2005; Tian et al., 2017), suggesting a crucial role of 13-*S*-HODE as a tumor suppressive metabolite. However, the effect of 13-*S*-HODE on mTOR signaling has not yet been profoundly investigated. Colorectal cancer (CRC) cell lines (HCT116, SW480, and HT29) were assessed to test the role of 13-*S*-HODE in cellular mTOR signaling events. Treatment of the CRC cells with 13-*S*-HODE for 48 h reduced phosphorylation at T389 of S6K1 and S240/244 of S6 but not S473 of Akt (Figure 4A), thereby demonstrating 13-S-HODE’s inhibitory effect on mTORC1. Similar mTORC1 inhibition was observed in other cancer cells such as MDA-MB-468 breast cancer cells and U-373 MG glioblastoma, as shown by the decreased phosphorylation of S6K1 and S6 upon treatment with 13-*S*-HODE (Figure 4B). In addition, the treatment of MDA-MB-468 cells with 13-*S*-HODE resulted in reduced proliferation (Figure 4C) and colony formation (Figure 4D). The effects of 13-*S*-HODE on tumor growth in vivo were examined by injecting MDA-MB-468 cells subcutaneously into immunocompromised mice and monitoring tumor growth. The growth of MDA-MB-468 xenografts was significantly inhibited by 13-*S*-HODE administration (Figures 4E and 4F), thus confirming its anti-cancer effects in vivo.

**Figure 4.**
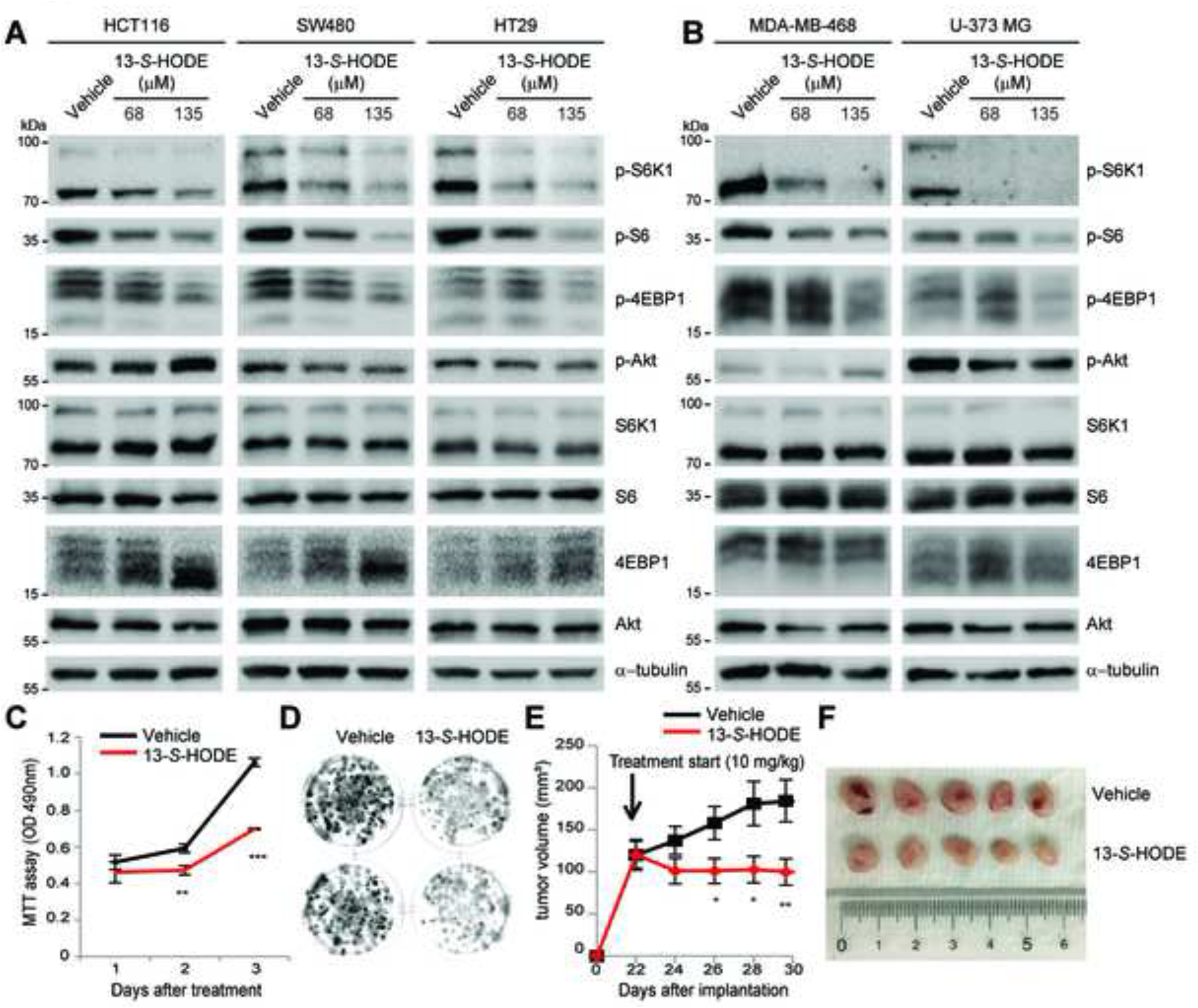
13-*S*-HODE inhibits mTOR signaling and cancer cell growth. (A) HCT116, SW480, and HT29 colorectal cancer cells were treated with DMSO vehicle or 13-*S*-HODE for 48 h. Cell lysates were analyzed by immunoblotting for the levels of the indicated proteins and phosphorylation states. (B) MDA-MB-468 breast cancer cells and U-373 MG glioblastoma cells were treated with DMSO vehicle or 13-*S*-HODE for 48 h. Cell lysates were analyzed by immunoblotting for the levels of the indicated proteins and phosphorylation states. (C and D) Cell viability (C) and colony formation (D) of DMSO vehicle or 135 μM 13-S-HODE treated MDA-MB-468 cells were measured. (E and F) MDA-MB-468 cells were inoculated into flanks of nude mice (*n* = 6 per group) and tumor volumes were measured for 30 days after injection. DMSO vehicle or 13-*S*-HODE (10 mg/kg) were intratumorally injected 5 times into xenograft tumors every 2 days (E). Data are expressed as mean ± SE (\**P* < 0.05; \*\**P* < 0.01; \*\*\**P* < 0.001, Student’s *t* test.). Representative images of xenograft tumors at the day of sacrifice (F).

### 13-*S*-HODE suppresses mTOR signaling and the growth of cancer cells with hyperactive *MTOR* mutation

Recently, gain-of-function *MTOR* mutations have been discovered from several types of cancers and serve as biomarkers to predict their responses to mTOR inhibitors (Grabiner et al., 2014). Based on our findings that 13-*S*-HODE inhibits mTOR, we put forward the hypothesis that 13-*S*-HODE could retard the growth of cancer cells with a hyperactive *MTOR* mutation. To examine this possibility, we selected JHUEM7 endometrial cancer cells harboring the mTORC1-activating *MTOR* mutation S2215Y (Grabiner et al., 2014); 13-*S*-HODE treatment resulted in decreased proliferation as well as colony formation of JHUEM7 cells (Figures 5A, 5B, and S3*A*). As it has been reported for other cancer cells (Moussalli et al., 2011; Shureiqi et al., 1999), endogenous 13-*S*-HODE was almost undetectable in vehicle treated JHUEM7 cells, whereas it was approximately 150 μM in JHUEM7 cells treated with 13-*S*-HODE for 48 h (Figure S3B), thereby confirming efficient intracellular uptake of 13-*S*-HODE. To investigate the effects of 13-*S*-HODE on global gene expression in JHUEM7 cells, we performed RNA-sequencing. 13-*S*-HODE treatment downregulated 3729 genes and up-regulated 3842 genes (Figure 5C). Specifically, according to KEGG pathway enrichment analyses (Figure S4A), genes in the mTOR signaling pathway were markedly altered by 13-*S*-HODE treatment, which supports its action in the mTOR pathway. Significant changes in gene sets linked to various cancer-related pathways further suggest 13-*S*-HODE’s anti-cancer activity (Figure S4).

**Figure 5.**
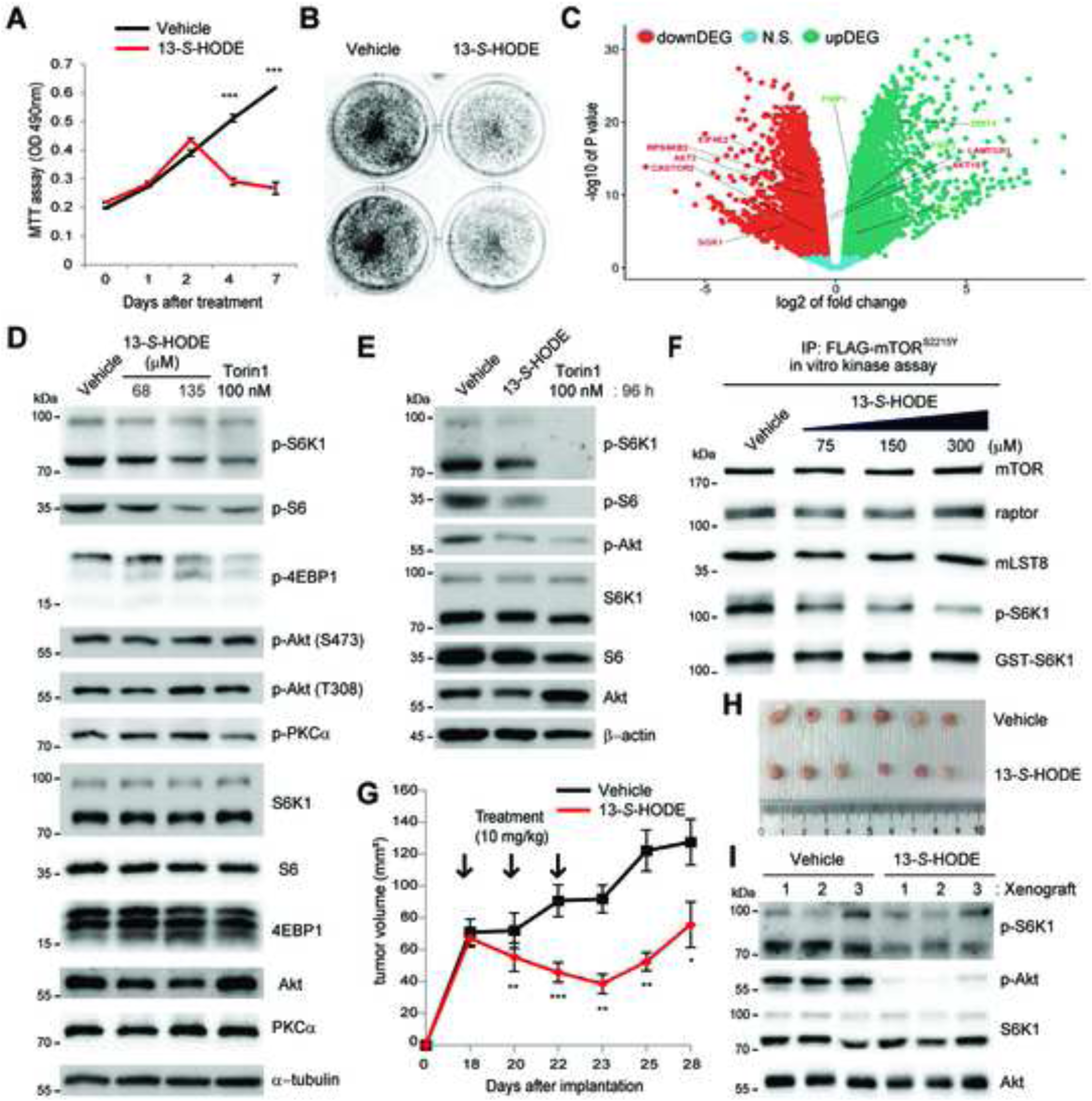
13-*S*-HODE inhibits mTOR signaling and suppresses the growth of cancer cells with hyperactive *MTOR* mutation. (A and B) Cell viability (A) and colony formation (B) of DMSO vehicle or 135 μM 13-*S*-HODE treated JHUEM7 endometrial cancer cells were measured. (C) Volcano plot showing significant gene expression changes in response to 135 μM 13-*S*-HODE treatment in JHUEM7 cells. (D) JHUEM7 cells were treated with DMSO vehicle or 13-*S*-HODE for 48 h. Cell lysates were analyzed by immunoblotting for the levels of the indicated proteins and phosphorylation states. (E) JHUEM7 cells were treated with DMSO vehicle or 135 μM 13-*S*-HODE for 96 h. Cell lysates were analyzed by immunoblotting for the levels of the indicated proteins and phosphorylation states. (F) In vitro mTORC1 activity of mTOR immunoprecipitates prepared from HEK293T cells expressing mTOR^S2215Y^ was assayed in the presence of 13-*S*-HODE. (G–I) JHUEM7 cancer cells were inoculated into flanks of nude mice (*n* = 6 per group) and tumor volumes were measured for 28 days after injection. DMSO vehicle or 13-*S*-HODE (10 mg/kg) were intratumorally injected 3 times into xenograft tumors every 2 days (G). Data are expressed as mean ± SE (\**P* < 0.05; \*\**P* < 0.01; \*\*\**P* < 0.001, Student’s *t* test.). Representative images of xenograft tumors at the day of sacrifice (H). Immunoblot analysis of tumor tissues harvested from xenograft (I). Data are representative of three independent experiments.

Next, we found that 13-*S*-HODE treatment reduced phosphorylation at T389 of S6K1 but not S473 of Akt (Figure 5D), thereby validating potent mTORC1 inhibition by 13-*S*-HODE. As shown in an mTOR kinase assay (Figures 2A and 2B), mTORC1 appears more sensitive to 13-S-HODE than mTORC2 (Figure 5D), presumably reflecting structural differences between the two distinct mTOR complexes (Aylett et al., 2016; Chen et al., 2018; Stuttfeld et al., 2018; Tafur et al., 2020; Yang et al., 2013; Yang et al., 2016). Interestingly, longer treatment of 13-*S*-HODE for 96 h decreased phosphorylation of both S6K1 T389 and Akt S473 (Figure 5E), suggesting that prolonged application of 13-*S*-HODE leads to the inhibition of both mTORC1 and mTORC2 in cells. In our immunoprecipitation mTOR kinase assays using the hyperactive mTOR mutants mTOR^S2115Y^, mTOR^C1483Y^, and mTOR^E1799K^, their kinase activities were dramatically suppressed in the presence of 13-*S*-HODE (Figures 5F and S5). Similarly, inhibition of hyperactive mTOR signaling and cellular growth was also found in 13-*S*-HODE–treated HEC59 endometrial carcinoma cells harboring the mTORC1-activating *MTOR* E1799K mutation (Figure S6). These results validate the suppressive effect of 13-*S*-HODE on cancer cells with a hyperactive *MTOR* mutation. Furthermore, the effect of 13-*S*-HODE on tumor growth in vivo was examined by injecting JHUEM7 cells subcutaneously into immunocompromised mice and monitoring tumor growth. mTOR signaling activities were markedly suppressed and growth of the JHUEM7 xenograft was stunted only when 13-*S*-HODE was injected intratumorally (Figures 5G–5I) or intravenously (Figure S7), thus confirming 13-*S*-HODE’s anti-tumorigenic potentials.

### ALOX15 overexpression reduces mTOR signaling and cancer cell growth

To further examine our hypothesis, we tested whether mTOR signaling can be affected by the expression of ALOX15 which is found down-regulated in many cancer cells (Moussalli et al., 2011; Shureiqi et al., 1999). We first chose MCF7 breast cancer cells with active PI3K signaling pathway. Compared with control cells expressing empty vector (MCF7^vector^), MCF7 cells stably overexpressing wild-type ALOX15 (MCF7^ALOX15 WT^) exhibited dramatically reduced phosphorylation of S6K1 T389 and Akt S473 (Figure 6A), indicating inhibition of both mTORC1 and C2. In contrast, no significant decrease in mTOR signaling was observed upon expression of catalytically-inactive mutant ALOX15 T560M (MCF7^ALOX15 T560M^) (Schurmann et al., 2011) (Figure 6A), suggesting the role of catalytic action of ALOX15 in the suppression of mTOR in cancer cells. Consistent with these observations, expression of ALOX15-WT, but not ALOX15-T560M, suppressed colony formation of MCF7 cells (Figure 6B). We also established hyperactive *MTOR*-harboring JHUEM7 cells stably overexpressing ALOX15-WT. Compared to vector-expressing control cells (JHUEM7^vector^), overexpression of ALOX15 in JHUEM7 cells (JHUEM7^ALOX15^) led to decreased phosphorylation of both S6K1 T389 and Akt S473 (Figure 6C), followed by reduced growth of cell culture (Figure 6D) and colony formation (Figure 6E) as well as tumor xenograft (Figures 6F and 6G). Collectively, these results clearly support a model in which 13-*S*-HODE is the key determinant of mTOR signaling activity and cell growth in cancer cells.

**Figure 6.**
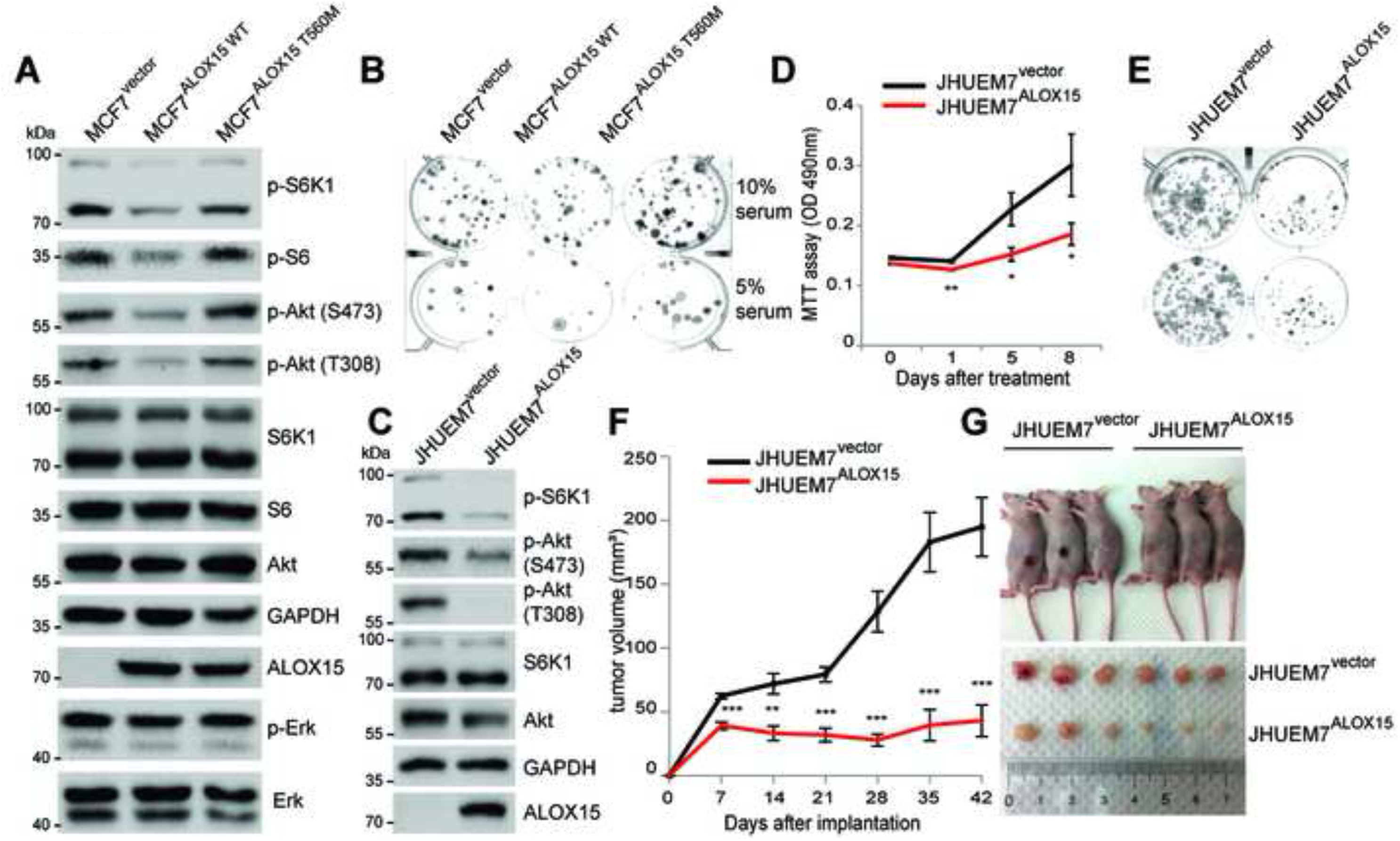
ALOX15 overexpression reduces mTOR signaling and cancer cell growth. (A and B) MCF7 cells with stable overexpression of empty vector, ALOX15-WT, or ALOX15-T560M were generated. Cell lysates were analyzed by immunoblotting for the levels of the indicated proteins and phosphorylation states (A). Colony formation of MCF7 cells in (*A*) was measured (B). (C–G) JHUEM7 cells with stable overexpression of empty vector or ALOX15-WT were generated. Cell lysates were analyzed by immunoblotting for the levels of the indicated proteins and phosphorylation states (C). Cell viability (D) and colony formation (E) of JHEUM7 cells were measured. JHUEM7 cells were inoculated into flanks of nude mice (*n* = 6 per group) and tumor volumes were measured for 42 days after injection (F). Data are expressed as mean ± SE (\**P* < 0.05; \*\**P* < 0.01; \*\*\**P* < 0.001, Student’s *t* test.). Representative images of xenograft tumors at the day of sacrifice (G).

## Discussion

Metabolite-protein interactions are key molecular events in mediating control of the fundamental biological processes between protein-based signaling networks and cellular metabolites (Li et al., 2010; Li and Snyder, 2011; McFedries et al., 2013; Tagore et al., 2008). Using an unbiased MS-based screening approach, we discovered 13-*S*-HODE as an mTOR-associated metabolite and identified its unique ability to inhibit the kinase activity of mTOR via directly interacting with its ATP-binding catalytic core. Furthermore, we demonstrated that 13-*S*-HODE supplementation or expression of ALOX15 in cancer cells suppresses mTOR signaling and attenuates cancer growth at both the cellular and physiological level.

mTOR functions as a signaling hub to integrate metabolic alterations such as amino acid uptake and increased glucose utilization (Kim and Guan, 2019; Saxton and Sabatini, 2017). Although information on the roles of amino acid- and glucose-sensing events in the regulation of mTOR signaling has emerged (Laplante and Sabatini, 2012; Orozco et al., 2020; Shimobayashi and Hall, 2014), our understanding of lipid metabolites in the coordination of mTOR has remained largely elusive. To the best of our knowledge, this is the first description of a molecular mechanism by which mTOR is negatively controlled by directly sensing a lipid metabolite. The range of 13-*S*-HODE actions toward mTOR in vitro and in cells was found to be in the higher micromolar level. Similarly, other cellular metabolites have shown to exert their biological effects on their protein targets in the micromolar up to the millimolar level, as exemplified from interactions such as fructose-1,6-bisphosphate with Sos1 (Son of sevenless homolog 1) (Peeters et al., 2017), lactate with NDRG3 (N-myc downstream-regulated gene 3) (Lee et al., 2015). In our biochemical characterization, we found that 13-*S*-HODE potently targets mTORC1 over mTORC2. Indeed, 48 h incubation of cancer cells with 13-*S*- HODE showed mTORC1 inhibition, but interestingly chronic treatment of 13-*S*-HODE over 96 h led to suppression of both the mTORC1 and C2 signaling pathways. 13-*S*- HODE’s action in the dual inhibition of mTORC1 and C2 was validated by robust reductions in both S6K1 and Akt signaling in 13-*S*-HODE−treated tumor xenograft tissues. This finding further highlights the value of 13-*S*-HODE’s signaling action because selective or incomplete inhibition of mTORC1 was reported to trigger feedback activation of the PI3K/Akt pathway (Benjamin et al., 2011; O’Reilly, 2006). Thus, further investigation is needed to understand signaling and molecular details underlying 13-*S*- HODE’s potent cellular impact on mTORC1/2 inhibition.

The levels of bioactive oxidized linoleic acid metabolites (OXLAMs) such as 13- S-HODE have been connected to the pathologies of major diseases such as cancer and coronary heart disease (Das, 2006; Vangaveti et al., 2016). Since mammals do not have enzymes to catalyze the de novo synthesis of linoleic acid, the intracellular extent to which it can be regulated (through diet, fatty acid transport, or metabolic transformation) needs to be uncovered. For instance, chronic headache and other pain sources can be managed in patients by controlling linoleic acid and its OXLAMs via dietary intervention (Ramsden et al., 2015). Modulation of fatty acid transport in cancer cells has also drawn attention because the control of fatty acid uptake in tumor cells is crucial for the progression of cancer (Corn et al., 2020; Watt et al., 2019). Approaches to overcome silenced ALOX15 expression in cancer cells might provide a way to employ 13-*S*-HODE mediated mTOR control. Thus, further investigations are required to understand fatty acid transporters and metabolic processes related to 13-*S*-HODE in cancer cells.

In summary, we uncovered a previously unknown metabolite-protein interaction that links PUFA metabolism in the case of the oxygenated linoleic acid metabolite, 13-*S*- HODE, to mTOR signaling in cancer. By expanding our classical view of 13-*S*-HODE as an endogenous ligand of the nuclear receptor PPAR (Itoh et al., 2008), we have established the suppressive action of 13-*S*-HODE on mTOR and cancer cell growth. Our findings suggest that epigenetic silencing of *ALOX15* expression and depletion of 13-*S*-HODE in multiple types of cancers have evolved to avoid its negative action on the mTOR signaling axis, which maintains mTOR activity and supports a favorable cellular milieu for tumor growth. Our study thus provides insight that targeting 13-*S*-HODE lipid metabolism in tumors might be an effective strategy to control mTOR in cancer cells.

## Significance

Mechanistic target of rapamycin (mTOR) is a key protein kinase that integrates various signals to control cell growth. Despite complex networks of mTOR with mTOR-associated proteins, endogenous cellular metabolites capable of interacting with mTOR have largely remained elusive. We identified that 13-*S*-hydroxyoctadecadienoic acid (13-*S*-HODE), a bioactive oxygenated metabolite of linoleic acid, a polyunsaturated essential fatty acid, directly binds to the catalytic domain of mTOR, thereby inhibiting its kinase activity. Either 13-*S*-HODE treatment or expression of arachidonate 15- lipoxygenase, an enzyme responsible for 13-*S*-HODE production, reduces mTOR and its downstream signaling activities, thus suppressing the growth of cancer cells as well as tumor xenografts. Thus, our data propose 13-*S*-HODE as a tumor suppressive, mTOR-inhibiting cellular metabolite involved in controlling cancer cell growth.

## Acknowledgements

We are grateful to Dr. Imad Shureiqi (University of Texas MD Anderson Cancer Center) for kindly providing ALOX15 expression plasmid. We also thank the members of the Byun and Kim laboratory, and Seung Min Park for their assistance and advice. This work was supported by the grant from the Samsung Science & Technology Foundation (SSTF-BA1401-14 to S.K.), TJ Park Science Fellowship of the POSCO TJ Park Foundation (to S.E.P.), National Research Foundation of Korea (NRF- 2020R1I1A1A01073144 to S.J.P.; NRF-2020R1A2C2005919 to Y.B.; NRF-2018R1A5A1024261, 2020R1A2C3005765 to S.K.), and KAIST Advanced Institute for Science-X fellowship (to S.J.P.). We also thank to Metabolomics core at the Convergence Medicine Research Center, Asan Medical Center for support and instrumentation.

## Author Contributions

S.J.P., Y.B., and S.K. conceived the work. S.J.P., H.M.K., A.Y., B.L., H.L., S.E.P., S.L., B.B.P., Y.K., J.L., and D.D. performed the experiments. S.J.P., H.M.K., B.L., B.O., D.D., P.L.W., M.Y.K., Y.B., S.K. designed the experiments and analyzed the data. S.J.P., H.M.K., D.D., Y.B. and S.K. wrote the manuscript.

## Methods

### LC-MS profiling of metabolites

Mass spectrometric analysis of the cell extracted samples was performed in a negative mode on a 1260 Infinity Quaternary Liquid Chromatography system equipped with a 6530 Q-TOF mass spectrometer (Agilent). Agilent MassHunter LC/MS Data Acquisition software (version B.05.00) was used for data collection. The mobile phases for liquid chromatography were water (A) and acetonitrile (B). Chromatography solvent system used was: 0-8 min (linear gradient): 0% solvent B to 100%, 8-10 min (isocratic): 100% solvent B. A 10 μL of sample dissolved in methanol was injected to a reversed-phase analytical column (Kinetex 2.6 μm XB-C18, 100 Å, 50 x 2.10 mm) with a flow rate of 0.5 mL/min. Typical sample ESI condition was set at nozzle voltage 1.0 kV, gas temperature 325 °C, drying gas 10 L/min, nebulizer 20 psig, VCap 4000 V, fragmentor 100 V and skimmer 65 V. All LC-MS data was deconvoluted into individual chemical peaks by the MassHunter Qualitative Analysis software B.05.00. The data files were transformed to CEF format files using DA reprocessor (Agilent, B.05.00). Subsequent statistical evaluation of independent pairs of groups were performed using Mass Profiler Professional software (Agilent, version 12.1) by unpaired t-test analysis (cut-off value of *P* < 1.0, cut-off value of fold change > 1.0). The features found in LC/MS data file was filtered based on an abundance filtering (minimum absolute abundance: 1000 counts) and number of ions (minimum number: 1). Percentile shift (value 75.0) was used as normalization algorithm option to compute the 75th percentile of the abundance values across all entities. ID Browser identification was conducted with a matching elemental formula by Agilent Masshunter ID Browser software B.05.00 using MassHunter METLIN Metabolite PCDL (Personal Compound Database and Library) file as a database.

### HDX-MS analysis

The protein sequence coverage map was obtained from undeuterated controls as follows: 3.5 μL of 50 μM mTOR C-terminal fragment in 25 mM HEPES, pH 7.4, 50 mM KCl, 10 mM MgCl_2_, 10% glycerol was diluted 96.5 μL of ice cold quench (100 mM Glycine, 25 mM TCEP, pH 2.5). The quenched samples were injected into a Waters HDX nanoAcquity UPLC (Waters) with in-line pepsin digestion (Waters Enzymate BEH pepsin column). Peptic fragments were trapped on an Acquity UPLC BEH C18 peptide trap and separated on an Acquity UPLC C18 column. A 7 min, 5% to 35% acetonitrile (0.1% formic acid) gradient was used to elute peptides directly into a Waters Synapt G2 mass spectrometer (Waters). MSE data were acquired with a 20 to 30 V ramp trap CE for high energy acquisition of product ions as well as continuous lock mass (Leu-Enk) for mass accuracy correction. Peptides were identified using the ProteinLynx Global Server 2.5.1 (PLGS) from Waters. Further filtering of 0.3 fragments per residues was applied in DynamX 3.0. HD exchange reactions, quenching and injection were performed using a using a LEAP autosampler controlled with HDxDirector. Briefly, 3.5 μL of ∼50 μM mTOR C-terminal fragment in 25 mM HEPES, pH 7.4, 50 mM KCl, 10 mM MgCl_2_, 10% glycerol, 1% DMSO with or without 130 μM 13- *S*-HODE (Cayman Chemical) was incubated in 31.5 μL of 25 mM HEPES, 99.99% D_2_O, pD 7.4, 50 mM KCl, 10 mM MgCl_2_, 1% DMSO with or without 130 μM 13-*S*-HODE. All reactions were performed at 25 °C. Prior to injection, deuteration reactions were quenched at various times (10 sec, 1 min, 10 min, and 2 h) with 65 μL of ice-cold 100 mM Glycine buffer, 5 mM TCEP pH 2.5. Back exchange correction was performed against fully deuterated controls acquired by incubating 3.5 μL of 50 μM of mTOR C-terminal fragment in 31.5 μL 25 mM HEPES, 99.99% D_2_O, pD 7.4, 50 mM KCl, 10 mM MgCl_2_, containing 6 M deuterated Guanidine DCl for 2 h at 25 °C prior to quenching. All deuteration time points were acquired in triplicates.

The deuterium uptake by the identified peptides through increasing deuteration time and for the fully deuterated control was determined using Water’s DynamX 3.0 software. The normalized percentage of deuterium uptake (%D) at an incubation time t for a given peptide was calculated as follows: %D = (100(m_t_-m_0_))/(m_f_-m_0_). With m_t_ the centroid mass at incubation time t, m_0_ the centroid mass of the undeuterated control, and m_f_ the centroid mass of the fully deuterated control. Percent deuteration difference plots (Δ%D) were generated using the percent deuteration calculated. Confidence intervals for the Δ%D plots were determined (Houde et al., 2011) and adjusted to percent deuteration using the fully deuterated controls. This approach involves the use of a two-criteria condition for determining the statistical significance of deuterium uptake differences observed for any given peptide: first, a difference in deuterium uptake at any single deuterium incubation time point (in colors) which is superior to the 98% confidence interval (dashed horizontal lines in Figures S2B and S2C) as determined using the overall standard deviation from the entire data set (all peptides, all time points, all states). Second, a summed difference in deuterium uptake integrated over all time points probed which is superior to its respective 98% confidence interval (dash-dot horizontal line in Figure S2C) as determined using the overall standard deviation propagated to the number of time point.

### Molecular modeling

The 13-*S*-HODE used for this study was created as a 2D structure by ChemBioDraw (ver. 11.0.1) and transferred to Chem3D Pro (ver. 11.0.1) to generate 3D structure. The structure of 13-*S*-HODE was saved as mol file format. The generated mol file transferred to Discovery Studio 3.0 (DS 3.0, Accelrys Inc., USA). The process of ligand preparation and optimization was performed by Prepare Ligands Module in DS 3.0. The PDB coordinates of human mTOR (PDB ID: 5FLC) were downloaded from the Protein Data Bank (24). Before conducting docking studies, water molecules were removed from the protein-ligand complexes and side chain bumps of amino acid residues were fixed. Hydrogen atoms were added by application of CHARMm Force Field, and Momany-Rone partial charge settings were used as default settings in DS 3.0. The potential binding site of 13-*S*-HODE in the ATP-biding site of mTOR was assigned to a spherical region with a radius of 20 Å from the key amino acids consisting of the ATP-binding site (Lys2187, Glu2190, His 2340, Asn2343, and Asp2357) (Yang et al., 2013). Docking studies of 13-*S*-HODE were performed using CDOCKER protocol implemented in DS 3.0 by modifying default settings slightly (top hits: 100, random conformations: 10, orientations to refine: 10, simulated annealing: true, Force field: CHARMm, Grid extension: 8.0, Ligand partial charge method: Momany-Rone). The best-docked pose of 13-*S*-HODE was analyzed and visualized by using the PyMOL software.

### Fluorescence-based thermal shift assay

The thermal shift assays were performed using the 7500 Real-Time PCR System (Applied Biosystems) melting curve program with a temperature increment of 1.0 °C and a temperature range of 25–95 °C. All reactions were incubated in a 20 μl final volume and assayed in 96-well plates using 1:1,000 dilution of 5,000 × SYPRO Orange stock solution (Sigma-Aldrich) and indicated concentrations (1.0 µM) of recombinant mTOR kinase domain diluted in buffer containing 10 mM HEPES·HCl pH 7.5. 13-*S*-HODE was added to the reaction to assess ligand-dependent thermal destabilization of mTOR kinase domain protein. The ligands (dissolved in DMSO) were incubated with mTOR kinase domain protein at 4 °C for 25 min before acquiring the melting curves (Huynh and Partch, 2015). The Tm is identified by plotting the first derivative of the fluorescence emission as a function of temperature (−dF/dT) using GraphPad PRISM7 software.

### Cell lines and tissue culture

HEK293 and HEK293T were cultured in high-glucose DMEM and 10% FBS (Atlas Biologicals) supplemented with 2 mM glutamine and 100 μg/ml penicillin/streptomycin. HCT116 (McCoy’s 5A), SW480,HT29 (RPMI), MDA-MB-468, MCF7, U373-MG (DMEM), and JHUEM7 (DMEM/F12 with 1x non-essential amino acid) cells were grown in each media supplemented with 10% FBS, 2 mM glutamine, and 100 μg/ml penicillin/streptomycin. All cell lines were maintained at 37 °C and 5% CO_2_. HEK293 and HEK293T were obtained from ATCC (American Type Culture Collection). JHUEM7 cells were obtained from RIKEN. All cell lines were validated and tested for mycoplasma.

### Generation of stable cell lines

The plasmid pCDH-MCS-T2A-copGFP-MSCV (System Biosciences; CD525A-1) was used as a backbone for expression of genes of interest. Lentiviruses were generated in HEK 293T cells by transfecting the lentiviral plasmids together with packaging vectors (pRSV-Rev, pMDLg/pRRE) and envelop expressing plasmid (pMD2.G) as manufacturer’s protocol. Lentiviruses were added to MCF7 cells for 24 h and the transduced cells were selected by flow cytometry.

### Cell lysis, immunoprecipitation, and immunoblotting

Cells were lysed in NP-40 lysis buffer (50 mM Tris-HCl at pH 7.5, 150 mM NaCl, 1% NP-40 substitute, 1 mM EDTA at pH 8.0, 50 mM NaF, 10 mM Na-pyrophosphate, 15 mM Na_3_VO_4_, and protease inhibitor cocktail (Roche)). Whole cell lysates were mixed with 5x SDS-sample buffer to a final concentration of 1-2 μg/μL, boiled for 5 min, and then used directly for immunoblotting or frozen at -20 °C. For mTOR immunoprecipitations, the cells were treated as with the glutathione pulldown, except they were lysed/washed in CHAPS buffer (40 mM HEPES at pH 7.4, 120 mM NaCl, 2 mM EDTA, 0.3% CHAPS, 50 mM NaF, 10 mM Na-pyrophosphate, and protease inhibitor cocktail (Roche)) and incubated with FLAG-M2-coupled agarose beads for 2 h at 4 °C. For immnunoblotting, 10-50 μg of whole cell lysates, or 10-30 μL of FLAG-M2 immunoprecipitates were loaded into lanes of 4-12% bis-tris SDS-PAGE gels, run at 120 volts for 2 h in 1x glycine buffer, and then transferred to 0.45 μm nitrocellulose membrane at 60 volts for 2 h in 1x transfer buffer (1x glycine buffer without SDS, 20% methanol). The membranes were then immunoblotted for the indicated proteins. All primary antibodies were diluted 1:1000 in 5% BSA W/V TBST, with the exception of GAPDH and FLAG antibodies which were diluted 1:2000. All secondary antibodies were diluted 1:5000 in 5% skim milk W/V TBST.

### In vitro mTOR kinase assay

For in vitro mTOR kinase assay in Figure 3F, HEK293 cells were used to immunoprecipitate endogenous mTOR complexes. For mTORC1 and C2 kinase assays in Figure 2A and 2B, FLAG-raptor or FLAG-rictor was cotransfected in HEK293T cells for 48 h with myc-tagged mTOR. In Figure S4, HEK293T cells were transfected with FLAG-mTOR^S2215Y^ for 48 h. Cells were rinsed once with ice-cold PBS and lysed in ice-cold CHAPS buffer. Cell lysates were incubated at 4 °C for 10 min and the supernatant was collected by centrifuging lysates at 13,000 rpm for 10 min. 2 μg of mTOR or FLAG antibody were added to the 2 mg of cell lysates and incubated with rotation for 2 h at 4 °C. 20 μl of agarose beads (Pierce) were added and the incubation continued for 1 hr. mTOR or FLAG immunoprecipitates were washed twice with the same lysis buffer and twice with kinase wash buffer (25 mM HEPES at pH 7.4, 20 mM potassium chloride, 1 mM magnesium chloride). mTORC1 kinase assays were performed for 15 min at 37 °C in a final volume of 15 μl of mTORC1 kinase buffer (25 mM HEPES at pH 7.4, 50 mM KCl, 10 mM MgCl_2_, 500 μM ATP) and 150 ng of S6K1 as a substrate. Reactions were stopped by the addition of 10 μl of sample buffer and boiling for 5 min and analyzed by SDS-PAGE and immunoblotting. In vitro mTORC2 kinase assay was performed by using mTORC2 kinase buffer (25 mM HEPES at pH 7.5, 100 mM potassium acetate, 1 mM MgCl_2_, 500 μM ATP) with 100 ng of Akt1 as a substrate.

### In vitro kinase assays for PI3KC3, DNA-PK, Erk

Kinase assays were performed by Reaction Biology Corporation. For in vitro PI3KC3 kinase assay, 50 μM PI:PS was used as a substrate and the ADP production was quantified by ADP-Glo luminescence detection. Both DNA-PK and Erk kinase reaction buffers include 20 mM Hepes (pH 7.5), 10 mM MgCl_2_, 1 mM EGTA, 0.02% Brij35, 0.02 mg/ml BSA, 0.1 mM Na_3_VO_4_, 2 mM DTT, and 1% DMSO. To measure DNA-PK activity, 20 μM peptide substrate [EPPLSQEAFADLWKK], 10 μg/ml DNA, and 10 μM ATP were used. For in vitro Erk kinase assay, 20 μM myelin basic protein and 10 μM ATP were used. ^33^P-ATP (specific activity 10 μCi/μl) into the reaction mixture to initiate the reaction. After 2 h of reaction at room temperature, radioactivity was measured by filter-binding method. Kinase activity data were expressed as the percent remaining kinase activity in test samples compared to vehicle (DMSO) reactions.

### Biotin-labeled 13-*S*-HODE pull-down assay

HEK293T cells were lysed in CHAPS lysis buffer used for mTORC immunoprecipitation. 2 mg of total cell lysates were incubated with 10 μM of biotinylated 13-*S*-HODE (Cayman Chemical) or linoleic acid (Sigma). For competition, 20 μM of non-biotinylated 13-*S*-HODE or linoleic acid were pre-incubated with cell lysates before binding. Streptavidin beads were added to the reaction mixture, and the assay was assessed by immunoblotting. To map 13-*S*-HODE–mTOR binding, HEK293T cells were transfected with the FLAG-mTOR^WT^, mTOR^D2195A^ or mTOR^K2187A/E2190A^ and then lysed in CHAPS lysis buffer. 2 mg of cell lysates were incubated with 20 μl of beads conjugated with an antibody against FLAG (Sigma-Aldrich) for 18 h at 4 °C with agitation. Then, beads were washed and incubated with or without unlabeled 13-*S*-HODE in the presence of biotinylated 13-*S*-HODE.

### Measurement of cellular 13-*S*-HODE by LC-MS/MS

13-*S*-HODE was extracted from ∼1.4 mL cell supernatant using solid phase extraction (SPE). 60 mg Oasis HLB (Waters) SPE cartridge was washed and preconditioned with ethylacetate, methanol, and (0.1% acetic acid + 5% MeOH) in H_2_O, sequentially. 10 μL of 0.2 mg/mL of EDTA and BHT in MeOH: H_2_O (50:50) was added to sorbent bed of a SPE column. Equal volume of (0.1% acetic acid + 5% MeOH) in H_2_O was added to the cell supernatant. 50 μL of 500 nM 13-*S*-HODE-d4 was also added as internal standard into a sample solution. The solution was added into SPE column. The column was washed with 2 column volumes of (0.1% acetic acid + 5% methanol) in H_2_O, then the column was dried using vacuum. Finally, 13-*S*-HODE was eluted with 0.5 mL methanol followed by 1.5 mL ethylacetate. The sample solution was dried using vacuum centrifuge, then stored at -20 °C until analysis. The dried matter was reconstituted with 40 μL of methanol prior to LC-MS/MS analysis.

13-*S*-HODE was determined using an LC-MS/MS system equipped with Agilent 1290 HPLC (Agilent) and QTRAP 5500 mass spectrometry (AB Sciex). A reverse-phase column (Pursuit5 C18, 150 × 2.1 mm) was used with mobile phase A (0.1% acetic acid in H_2_O) and mobile phase B (0.1% acetic acid in acetonitrile/methanol (84/16)). The LC was run at 250 μl/min and 35 °C. The separation gradient was as follows: 35% B at 0 min, hold at 35% B for 0.25 min, 35 to 45% of B for 0.75 min, hold at 45% of B for 2 min, 45 to 56% of B for 5.5 min, 56 to 65% of B for 5.5 min, hold at 65% of B for 6 min, 65 to 95% of B for 1.5 min, hold at 95% of B for 5.5 min, 95 to 35% of B for 0.1 min, and then hold at 35% of B for 2.9 min. The multiple reaction monitoring (MRM) mode was performed in negative ion mode for 13-*S*-HODE, and the extracted ion chromatogram corresponding to the specific transition of 13-*S*-HODE was used for quantification (Q1/Q3= 295/195, RT = 18.09 min for 13-*S*-HODE; Q1/Q3= 299/198, RT = 17.93 min for 13-*S*-HODE-d4). The calibration range for 13-HODE was 1-10000 nM (r2 ≥ 0.99). Data analysis was performed by using either Analyst 1.5.2.

### Cell proliferation and colony formation assay

MDA-MB-468, JHUEM7 (5 × 10^3^ cells/ml), MDA-MB-231, MCF7 (1 × 10^4^ cells/ml) were seeded into 96-well culture plates. After overnight incubation, cells were treated with DMSO or 135 μM of 13-*S*-HODE for various times. Then 20 ml 3-(4,5- dimethylthiazol-2-yl)-2,5-diphenyl tetrazolium bromide (MTT) solution (2.5 mg/ml in PBS) was added to each well, and the plates were incubated for an additional 1 hr at 37°C. The optical density of each well was measured at 492 nm with a SpectraMax Paradigm Reader (Molecular Devices). For colony formation assay, cells (4 × 10^4^/well) were seeded into 24-well plates and cultured overnight before being incubated with DMSO or 135 μM of 13-*S*-HODE for 4 days. The resulting colonies were fixed with methanol at −20 °C for 30 min and were stained with 1% crystal violet for 15 min.

### RNA-sequencing

Total RNA was isolated from cells using Tri reagent (Molecular Research Center) according to the manufacturer’s protocol. 1 μg of total RNA was processed for preparing mRNA sequencing library using MGIEasy RNA Directional Library Prep Kit (MGI) according to manufacturer’s instruction. The first step involves purifying the poly-A containing mRNA molecules using poly-T oligo attached magnetic beads. Following purification, the mRNA is fragmented into small pieces using divalent cations under elevated temperature. The cleaved RNA fragments are copied into first strand cDNA using reverse transcriptase and random primers. Strand specificity is achieved in the RT directional buffer, followed by second strand cDNA synthesis. These cDNA fragments then have the addition of a single ’A’ base and subsequent ligation of the adapter. The products are then purified and enriched with PCR to create the final cDNA library. The double stranded library is quantified using QauntiFluor ONE dsDNA System (Promega). The library is circularized at 37 °C for 30 min, and then digested at 37 °C for 30 min, followed by cleanup of circularization product. To make DNA nanoball (DNB), the library is incubated at 30 °C for 25 min using DNB enzyme. Finally, library was quantified by QauntiFluor ssDNA System (Promega). Sequencing of the prepared DNB was conducted on the MGIseq system (MGI) with 150 bp paired-end reads. The limma, edgeR, msigdbr, clusterProfiler packages in R, an open-source programming environment, was used to perform differentially expressed genes, gene set enrichment and pathway enrichment analysis. ENTREZID, MsigDB, GO terms and KEGG pathways were mapped and were used to perform enrichment test based on hypergeometric distribution. To prevent high false discovery rate (FDR) in multiple testing, q-values were also estimated for FDR control.

### Mouse tumor xenograft studies

Animal protocols were performed in accordance with the guidelines approved by the Korea Advanced Institute of Science and Technology Animal Care and Use Committee. MDA-MB-468 (1 × 10^7^ cells), JHUEM7 (5 × 10^6^ cells) cells were implanted subcutaneously into the female BALB/c nude mice at the age of 8-12 weeks. After the mean tumor volumes reached 50 mm^3^, the mice were randomly assigned into two different groups (6 mice/group). The body weight and tumor diameters were measured every other day. Tumor volume was assessed using calipers according to the formula (0.5) x (W)^2^ x (L), and a student’s t test was used to determine the *P* values. Mice received intratumoral or intravenous injections of 10 mg/kg 13-*S*-HODE every 2 days.

## Data availability

All data needed to evaluate the conclusions in the paper are present in the paper and/or the Supplemental information. All other data are available from the corresponding author upon reasonable request.

## Competing interests

The authors declare no competing interests.

## Notes

### Competing Interest Statement

The authors have declared no competing interest.

