## Supplemental Information for "Linoleic acid metabolite, 13-*S*-hydroxyoctadecadienoic acid, suppresses cancer cell growth by inhibiting mTOR"

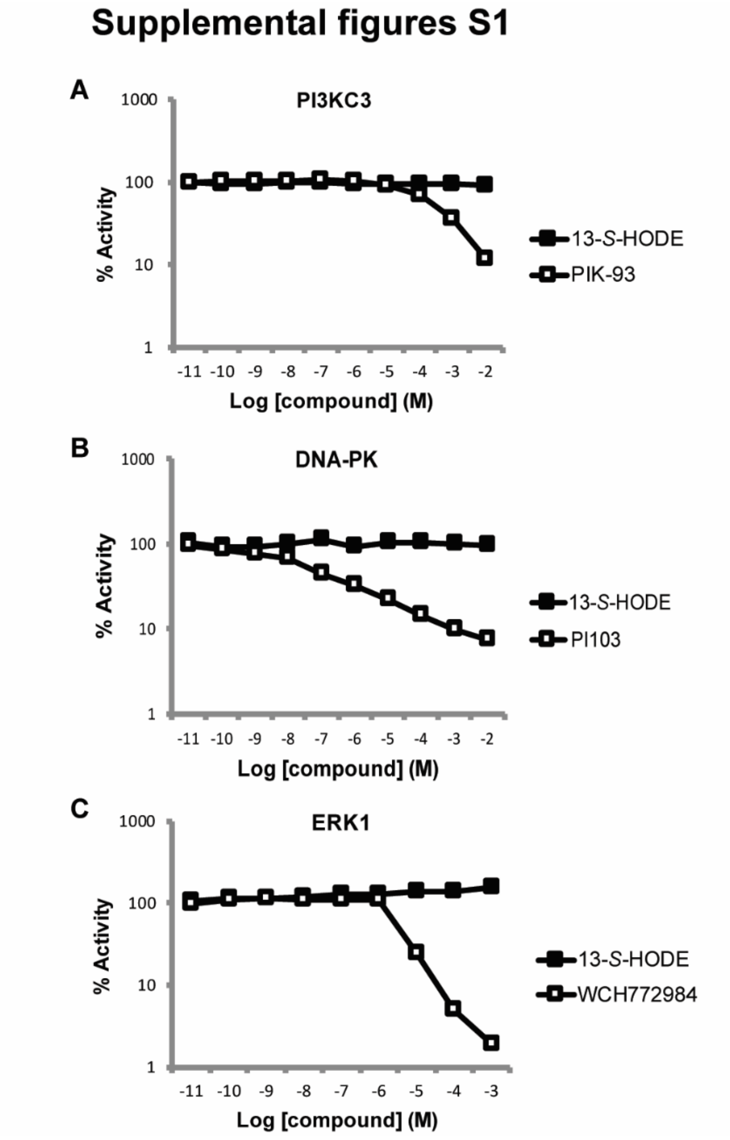


**Fig. S1. 13-*S*-HODE is selective for mTOR over related kinases.**
(A–C) In vitro kinase assays were performed using the recombinant human phosphatidylinositol 3-kinase catalytic subunit type 3 (PI3KC3) (A), DNA-dependent protein kinase catalytic subunit (DNA-PK) (B), or ERK (C). PIK-93, PI103, WCH772984 were used for positive control inhibitors for PI3KC3, DNA-PK, ERK, respectively.

### Supplemental figures S2-1

**A**

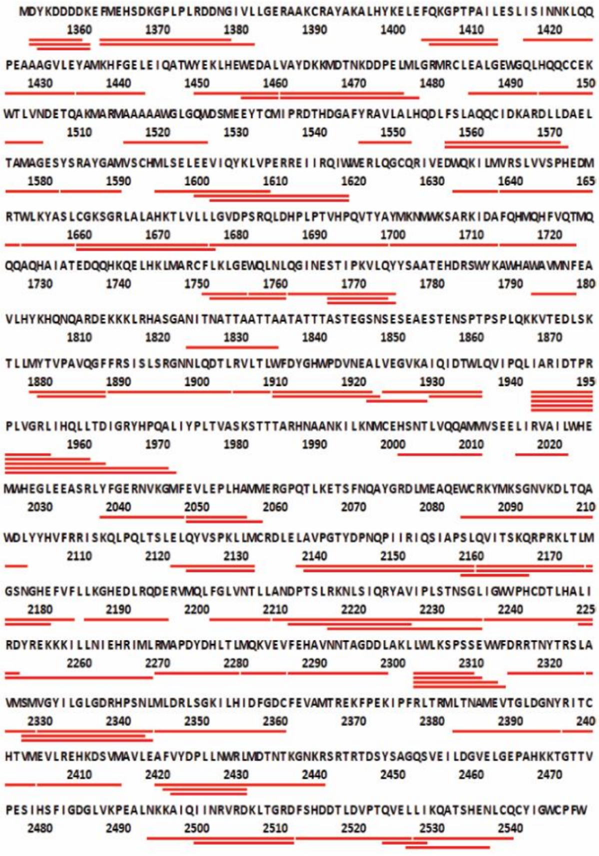

Total: 100 peptides, 66.7% Coverage, 1.61 Redundancy

**B**

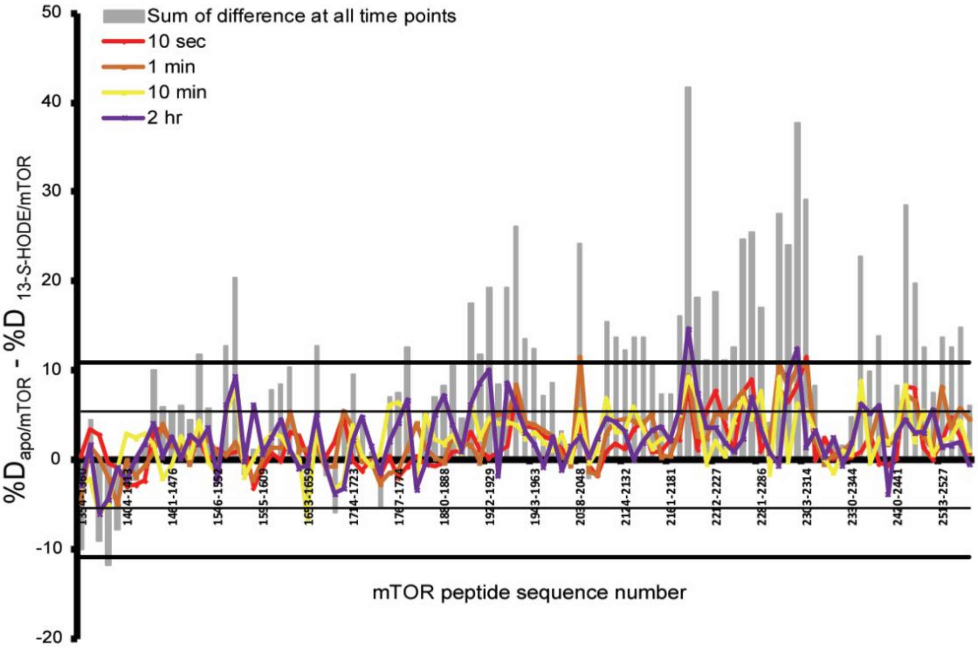

### Supplemental figures S2-2 (continued)

C

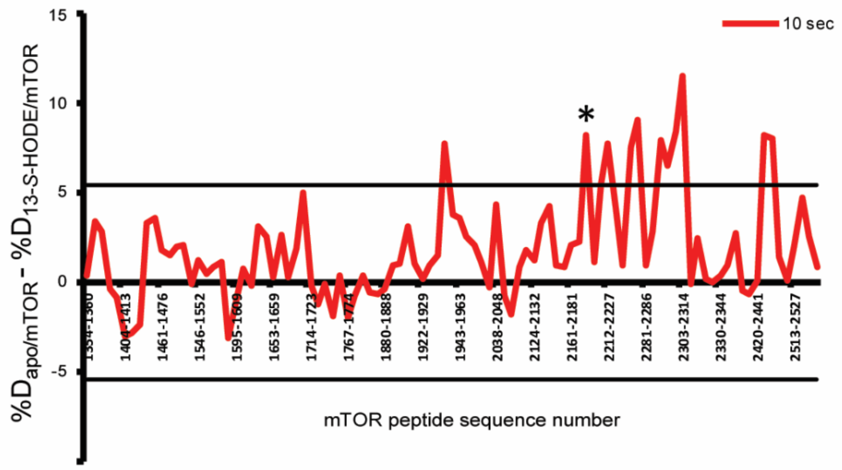


D

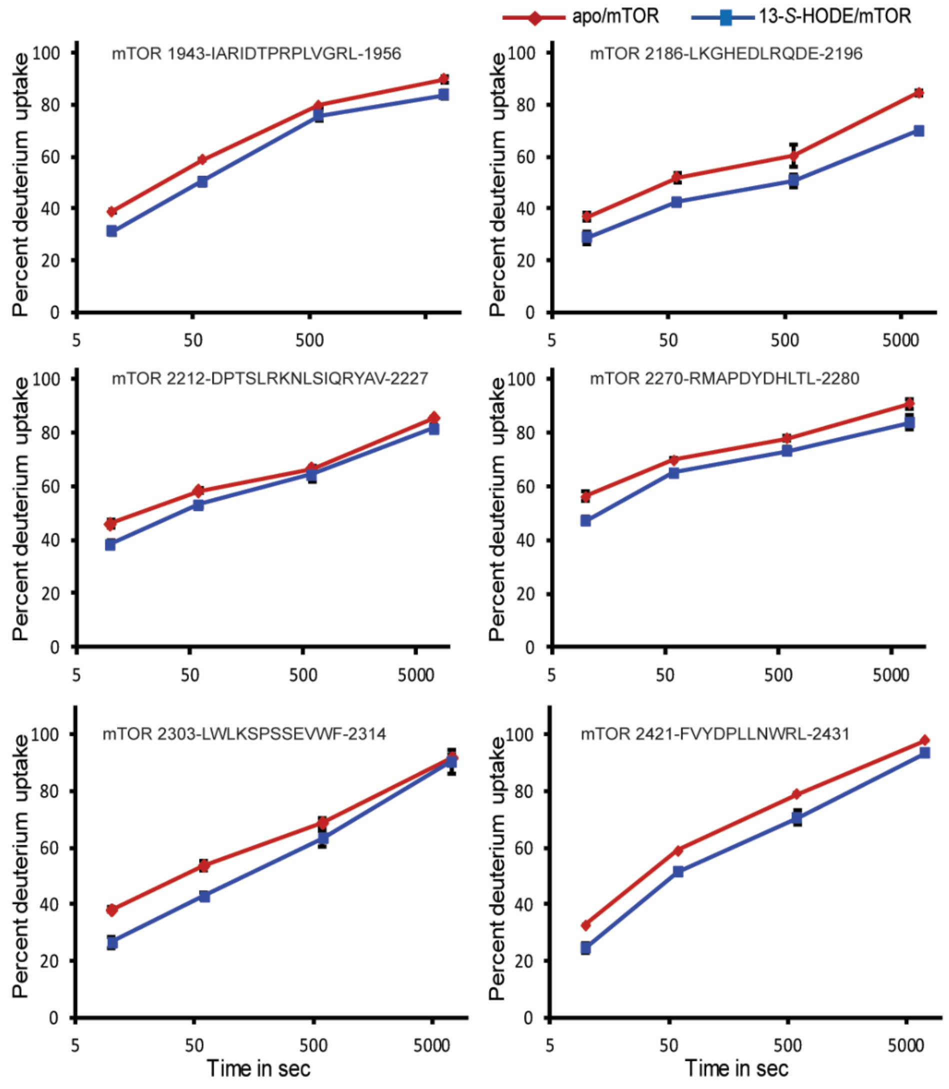

**Fig. S2. HDX-MS analysis of 13-S-HODE-mTOR C-ter.**

(A) Sequence coverage map of mTOR obtained for HDX-MS. Peptides that were successfully identified and of sufficient quality for HDX-MS are displayed as red bars under the sequence.

(B) Difference plots ($\%D_{apo/mTOR}$ - $\%D_{13\text{-}S\text{-}HODE/mTOR}$) for all peptides at all time point. Individual peptides are listed on the x-axis from N to C terminus. Deuterium labelling time points are color coded according to the legend. Grey bars represent the sum of the difference in %D integrated over all time points. Respective 98% confident intervals are indicated by solid lines.

(C) Difference plots ($\%D_{apo/mTOR}$ - $\%D_{13\text{-}S\text{-}HODE/mTOR}$) for all peptides at 10 sec of deuterium labeling. 98% confident intervals are indicated by solid lines. Peptides showing a difference superior to the 98% confident intervals are deemed statistically significant deviations.

(D) Deuterium uptake kinetic traces of individual six peptides detected in (C).

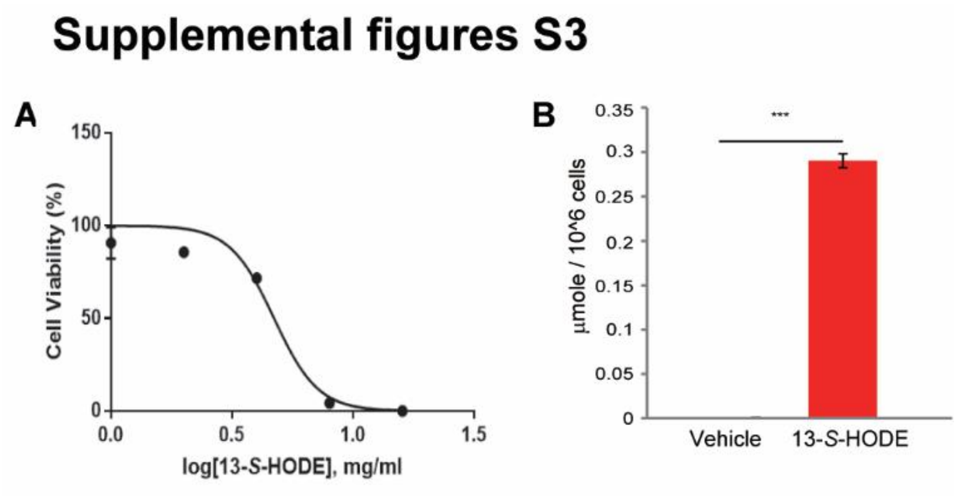


**Fig. S3. Inhibition of cancer cell growth by 13-*S*-HODE.**
(A) Cell viability was measured after treating JHEUM7 cells with DMSO vehicle or different concentrations of 13-S-HODE for 48 h. EC50 = ~4.7 mg/ml (~159 µM)
(B) Intracellular concentration of 13-*S*-HODE was measured from JHEUM7 cells with DMSO vehicle or 150 µM 13-*S*-HODE for 48 h. Data are expressed as mean ± SE (***$P < 0.001$).

### Supplemental figures S4

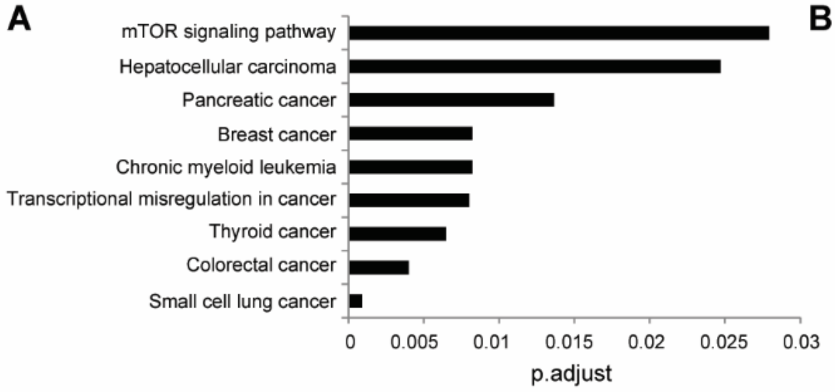


B

| KEGG pathways | p.adjust |
|---|---|
| Cell cycle | 5.24E-06 |
| Cellular senescence | 5.24E-06 |
| Apoptosis | 0.000252666 |
| Biosynthesis of amino acids | 0.004000862 |
| Ferroptosis | 0.008240041 |
| Longevity regulating pathway | 0.019330016 |
| Biosynthesis of unsaturated fatty acids | 0.029270933 |
| Autophagy - animal | 0.032042458 |
| Fatty acid metabolism | 0.045474304 |

**Fig. S4. RNA-sequencing reveals gene expression changes in 13-*S*-HODE-treated JHUEM7 cells.**

Histogram showing the significant GO terms (related to cancer (A) and biological processes (B)) of differentially-expressed genes in 135 μM 13-*S*-HODE–treated JHUEM7 cells, compared to vehicle-treated cells.

### Supplemental figures S5

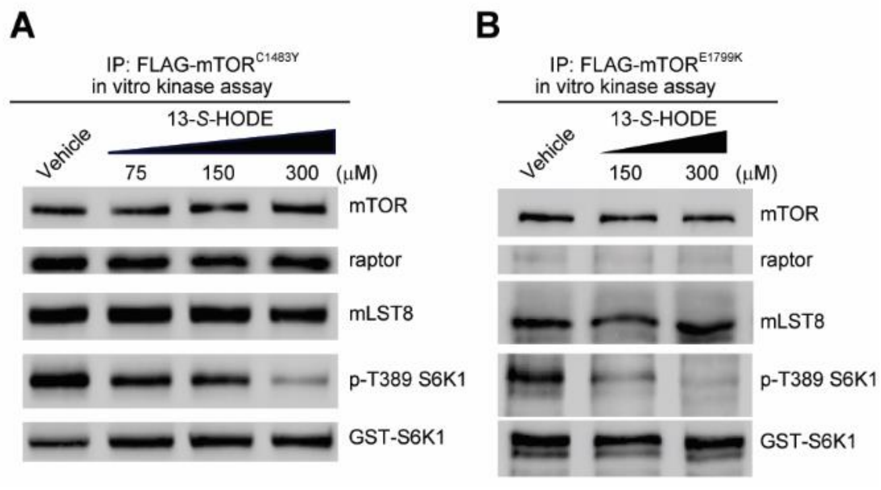


**Fig. S5. 13-*S*-HODE inhibits hyperactive mTOR mutants.**
(A and B) In vitro mTORC1 activity of mTOR immunoprecipitates prepared from HEK293T cells expressing mTOR$^{C1483Y}$ (A) or mTOR$^{E1799K}$ (B) was assayed in the presence of 13-*S*-HODE. S6K1 phosphorylation at T389 was evaluated via immunoblotting.

### Supplemental figures S6

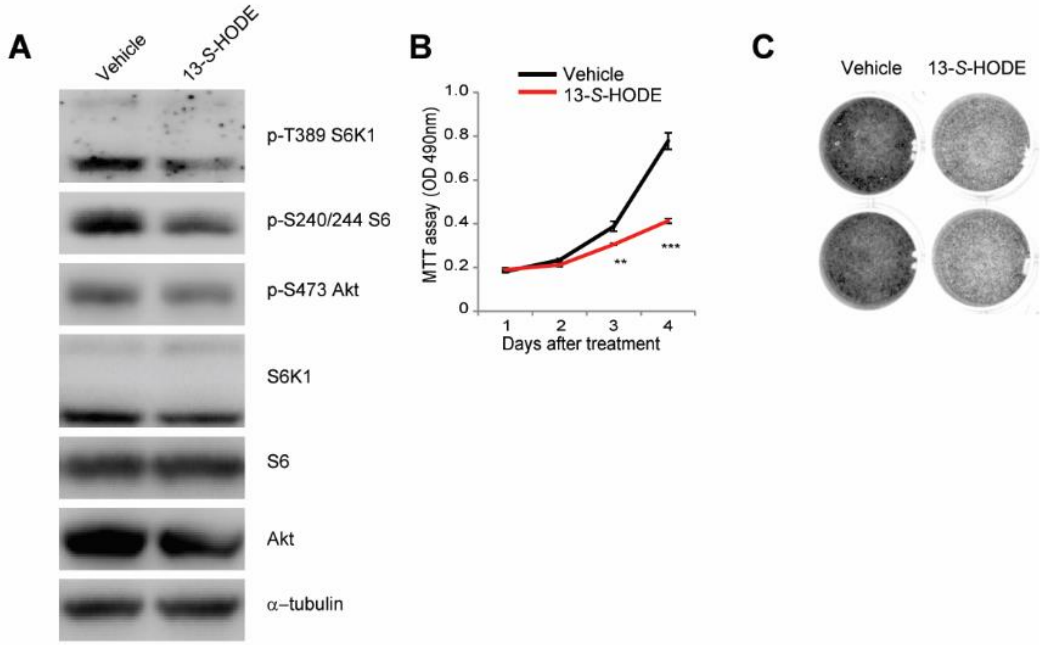


**Fig. S6. 13-*S*-HODE inhibits mTOR signaling and suppresses the growth of cancer cells with hyperactive *MTOR* mutation.**

(A) HEC59 cells harboring $mTOR^{E1799K}$ were treated with DMSO vehicle or 135 μM 13-*S*-HODE for 48 h. Cell lysates were analyzed by immunoblotting for the levels of the indicated proteins and phosphorylation states.

(B and C) Cell viability (B) and colony formation (C) of DMSO vehicle or 135 μM 13-*S*-HODE treated HEC59 endometrial cancer cells were measured.

### Supplemental figures S7

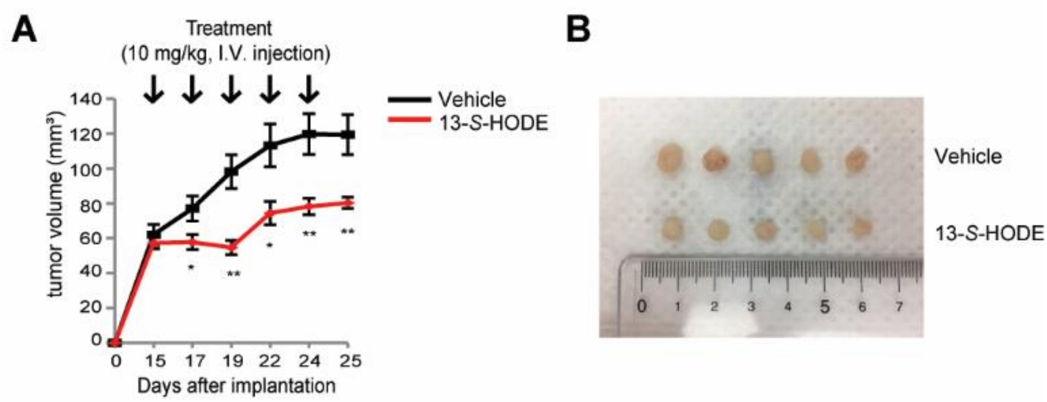


**Fig. S7. Intravenous injection of 13-*S*-HODE inhibits tumor growth in vivo.** (A and B) Nude mice bearing JHUEM7 xenografts were intravenously treated with 13-*S*-HODE (10 mg/kg) or vehicle. Tumor volumes were calculated and presented as growth curves (A). Data are expressed as mean ± SE (**P* < 0.05 and ***P* < 0.01). Representative tumor tissues resected from mice on day 10 after treatment (B).
